# *rab-27* acts in an intestinal secretory pathway to inhibit axon regeneration in *C. elegans*

**DOI:** 10.1101/2020.09.05.283267

**Authors:** Alexander T. Lin-Moore, Motunrayo J. Oyeyemi, Marc Hammarlund

## Abstract

Injured axons must regenerate to restore nervous system function, and regeneration is regulated in part by external factors from non-neuronal tissues. Many of these extrinsic factors act in the immediate cellular environment of the axon to promote or restrict regeneration, but the existence of long-distance signals regulating axon regeneration has not been clear. Here we show that the Rab GTPase *rab-27* inhibits regeneration of GABAergic motor neurons in *C. elegans* through activity in the intestine. Re-expression of RAB-27, but not the closely related RAB-3, in the intestine of *rab-27* mutant animals is sufficient to rescue normal regeneration. Several additional components of an intestinal neuropeptide secretion pathway also inhibit axon regeneration, including NPDC1/*cab-1*, SNAP25/*aex-4*, and KPC3/*aex-5*. Together these data indicate that RAB-27-dependent neuropeptide secretion from the intestine inhibits axon regeneration, and point to distal tissues as potent extrinsic regulators of regeneration.

## INTRODUCTION

Unlike many other tissues, where cells respond to injury through proliferation and replacement, cells in the nervous system are not usually replaced following axon damage. Instead, neurons rely on axon regeneration to restore the connectivity necessary for function. Despite its importance, however, axon regeneration is often inhibited *in vivo*, leading to permanent loss of nervous system function after injury.

A neuron’s axon regeneration capacity is extensively regulated by contacts with the extracellular environment of the injured axon. In the mammalian central nervous system, myelin-associated transmembrane signals Nogo, MAG and OMgp potently inhibit post-injury growth through direct interaction with neuronal receptors like Ngr1 and PTPσ (Liu et al. 2006, Cheah & Andrews 2016). In *C. elegans*, which lacks myelin-associated regeneration inhibitors, the peroxidasin PXN-2 and syndecan (SDN-1) control the integrity and signaling topography of the extracellular matrix to negatively or positively regulate regeneration success, respectively (Gotenstein et al. 2010, Edwards & Hammarlund 2014). Thus, a neuron’s local environment and neighbor cells influence its regenerative capacity.

In addition to responding to their local environment and neighbors, neurons respond to secreted, long-range signals from distant tissues, which can regulate neuronal programs ranging from synapse patterning to complex behaviors (Klassen & Shen 2007, Sawa & Korswagen 2013, Holzer & Farzi 2014). But for axon regeneration, the existence of long-range inhibitory signals *in vivo* has not been clear. We have previously identified the Rab GTPase *rab-27* as a conserved inhibitor of axon regeneration (Sekine et al. 2018). Here we show that *rab-27* inhibits regeneration of D-type motor neurons in *C. elegans* through activity in the intestine. We further show that inhibition of axon regeneration involves an intestinal secretory pathway involved in neuropeptide secretion. Together these results indicate that the *C. elegans* intestine inhibits axon regeneration, and point to long-distance, extrinsic signaling as a novel mechanism of axon regeneration regulation.

## RESULTS

### An intestinal function for RAB-27 in axon regeneration

*C. elegans* provides a robust system to investigate in vivo axon regeneration at single-neuron resolution (Hammarlund & Jin 2014). Previously, Rab27 was identified in a large-scale screen as a key inhibitor of regeneration (Sekine et al. 2018). This work demonstrated that Rab27B/*rab-27* inhibits regeneration in both mouse and *C. elegans* models, and indicated that one site of function for RAB-27 in *C. elegans* is in the injured neurons. However, in *C. elegans, rab-27* is highly expressed in the anterior- and posterior-most cells of the intestine as well as the nervous system (Mahoney et al. 2006, Cao et al. 2017). A potential function of *rab-27* in the intestine was not previously tested.

To study *rab-27*’s function in axon regeneration, we used the same regeneration assay as described in previous work (Sekine et al. 2018). We used the GABAergic neurons as our model system, lesioning individual axons with a pulsed laser and measuring subsequent regeneration (Fig. 1A). As shown previously, loss of *rab-27* resulted in high regeneration, with significant regeneration enhancement occurring as early as 12 hours after axotomy (Fig. 1B). *rab-27* mutants produced growth cones earlier and at a higher proportion than in controls, and axons of *rab-27* mutant animals that initiated regeneration grew further and reached the dorsal nerve cord earlier compared to control axons (Fig. 1C,D).

**Figure 1.**
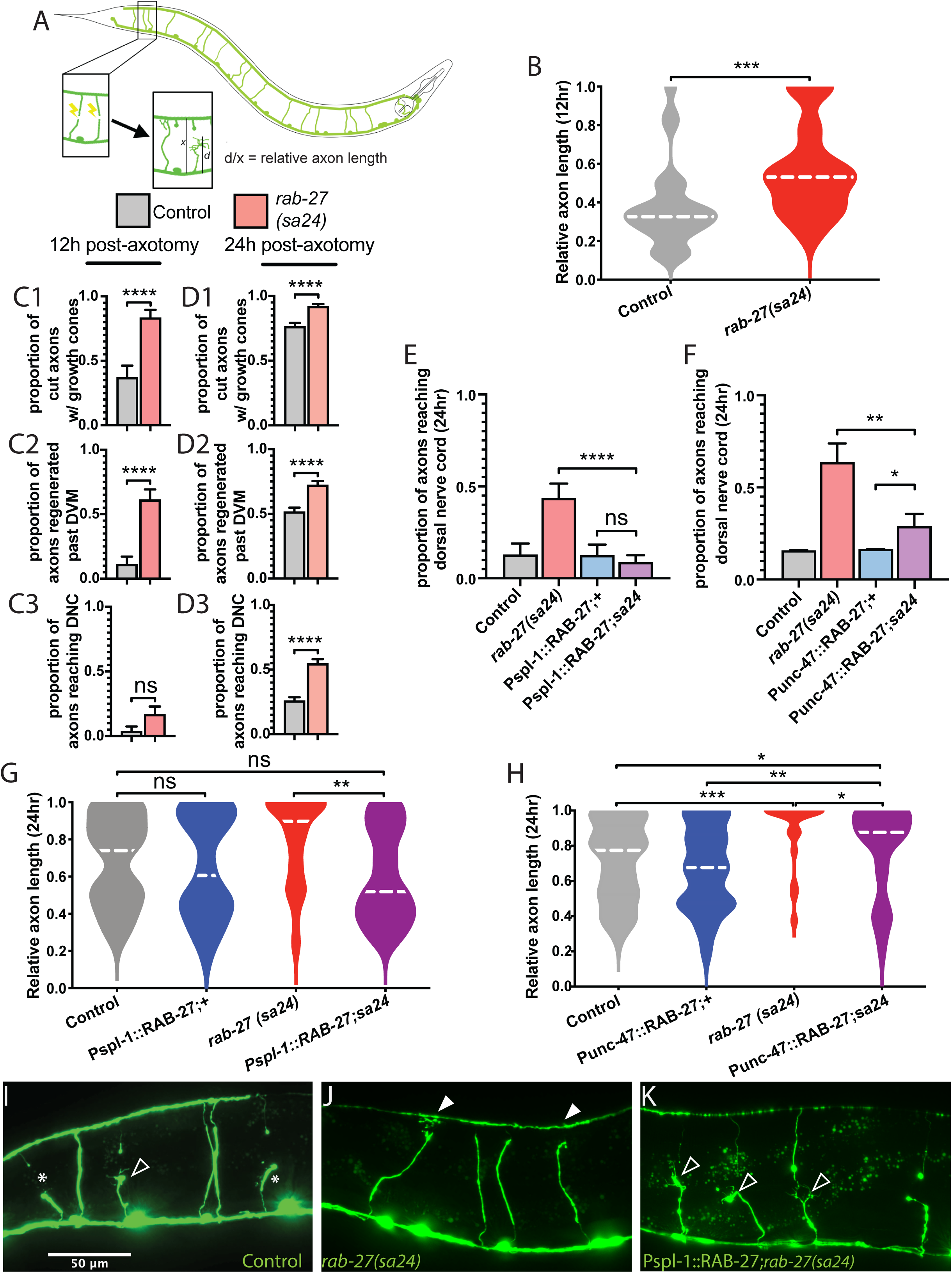
RAB-27 expression in the intestine inhibits axon regeneration. (A) Posterior DD/VD commissural axons in the GABAergic nervous system of L4 animals were severed using a pulsed laser, and regeneration was measured after a 24 hour recovery window. (B) Relative axon length in control (*oxIs12*) animals and *rab-27(sa24)* mutants after 12 hours of recovery after axotomy. Axons cut per genotype, L to R: 27, 36. Kolmogorov-Smirnov test was used. ns, not significant, * p < 0.05, *** p < 0.0005. (C). Proportion of cut axons forming growth cones (C1), regeneration past the dorsoventral midline (DVM) (C2), or full regeneration back to the dorsal nerve cord (DNC) (C3) in control (*oxIs12*) and *rab-27(sa24)* mutant animals after 12 hours of recovery post-axotomy. Axons cut per genotype, L to R: 27, 36. Unpaired t-test was used. ns, not significant, **** p < 0.0001. Error bars represent SEM. (D). Proportion of cut axons forming growth cones (D1), regeneration past the dorsoventral midline (DVM) (D2), or full regeneration back to the dorsal nerve cord (DNC) (D3) in control (*oxIs12*) and *rab-27(sa24)* mutant animals after 24 hours of recovery post-axotomy. Axons cut per genotype, L to R: 233, 198. Unpaired t-test was used. ns, not significant, **** p < 0.0001. Error bars represent SEM. (E) Proportion of cut axons showing signs of regeneration in control (*oxIs12*) and *rab-27(sa24)* mutant animals, and animals expressing RAB-27 cDNA under an intestine-specific promoter (Pspl-1) and stabilized with *rab-3* 3’ UTR sequence, in both control and *rab-27* mutant backgrounds. Axons were scored after 24 hours of recovery post-axotomy. Axons cut per genotype, L to R: 31, 39, 32, 57. Unpaired t-test was used. ns, not significant, **** p < 0.0001. Error bars represent SEM. (F) Proportion of cut axons showing signs of regeneration in control (*oxIs12*) and *rab-27(sa24)* mutant animals, and animals expressing RAB-27 cDNA under a GABA neuron-specific promoter (Punc-47) and stabilized with *rab-3* 3’ UTR sequence, in both control and *rab-27* mutant backgrounds. Axons were scored after 24 hours of recovery post-axotomy. Axons cut per genotype, L to R: 51, 22, 67, 45. Unpaired t-test was used. ns, not significant, * p < 0.05, ** p < 0.005. Error bars represent SEM. (G) Relative axon length in control (*oxIs12*) animals, *rab-27(sa24)* mutants, and animals expressing RAB-27 cDNA under an intestine-specific promoter and stabilized with *rab-3* 3’ UTR sequence, in both control and *rab-27* mutant backgrounds. Number of axons cut per genotype, L to R: 31, 32, 39, 57. Kolmogorov-Smirnov test was used. ns, not significant, * p < 0.05, ** p < 0.005. (H) Relative axon length in animals expressing RAB-27 cDNA under a GABA neuron-specific promoter, in both control (*oxIs12*) and *rab-27* mutant backgrounds. Number of axons cut per genotype, L to R: 51, 67, 22, 45. Kolmogorov-Smirnov test was used. ns, not significant, * p < 0.05, ** p < 0.005, *** p < 0.0005. (I-K). Representative micrographs of regeneration in Day 1 adults 24 hours after axotomy in *oxIs12* control (I), *rab-27* mutant (J), and intestinal *rab-27* rescue (K) animals. Filled arrows indicate fully regenerated axons reaching the dorsal nerve cord, empty arrows indicate partially regenerated axons, and stars indicate non-regenerating axon stumps. All animals express *Punc-47::GFP* (*oxIs12*).

Next, to determine whether intestinal *rab-27* might function in regeneration, we expressed *rab-27* in either the intestine or the neurons of mutant animals. The intestine is known to signal to the *C. elegans* nervous system to regulate the defecation motor program (Thomas 1990, Mahoney et al. 2008, Wang et al. 2013). However, signals from the intestine, which must travel through the pseudocoelom to reach the GABAergic neurons, have not previously been implicated in regulation of axon regeneration. We expected that expression in a tissue where it functions would restore normal, lower levels of regeneration. Surprisingly, re-expression of *rab-27* in the intestine of mutants was sufficient to significantly reduce regeneration compared to *rab-27* mutant animals (Fig. 1E, G, I-K), indicating that the intestine is a major site of *rab-27* function in inhibiting axon regeneration. Expression of *rab-27* in the GABA neurons of *rab-27* mutants also reduced regeneration relative to *rab-27* mutant animals, as previously described (Sekine et al. 2018). Thus, *rab-27* can function in both the intestine and in GABA neurons to inhibit axon regeneration.

Expression of *rab-27* in GABA neurons had a significant effect on regeneration but was not sufficient to fully suppress regeneration to control levels (Fig. 1F, Fig. S1A). By contrast, we previously found that expressing *rab-27* in GABA neurons restores regeneration to control levels (Sekine et al. 2018). Our current strategy to express *rab-27* only in GABA neurons used an expression construct that contained the *rab-3* 3’UTR, while our previous efforts used the *unc-54* 3’UTR. The *unc-54* UTR sequence can itself drive expression in the posterior gut because it contains regulatory and coding sequence for the intestinal gene *aex-5* (Silva-García et al. 2019). We hypothesized that a requirement for intestinal expression accounts for the different effects of the UTR. Intestine-specific *rab-27* rescue constructs containing the *rab-3* 3’UTR rescued axon regeneration identically to those containing the *unc-54* 3’UTR (Fig. S1B). Use of the *rab-3* 3’ UTR in the intestine-specific RAB-27 rescue construct also produced a much stronger rescue of *rab-27* mutants’ *aex* phenotype, with nearly full restoration of the pBoc/expulsion ratio, compared to only a partial rescue by constructs containing the *unc-54* 3’ UTR (Fig. S2). Thus, *rab-27* can act in either neurons or the intestine to suppress regeneration, but intestinal expression is necessary for complete function. Overall, these tissue-specific experiments raise the question of whether similar or different cellular mechanisms mediate *rab-27*’s regeneration function in these two tissues.

### RAB-27’s synaptic vesicle tethering cofactors do not inhibit regeneration

In neurons, *rab-27* is thought to function similar to the well-studied Rab family member *rab-3*. Phylogenetic analysis of the *C. elegans* Rab family shows that *rab-27* and *rab-3* are each other’s closest paralog (Gallegos et al. 2012). RAB-3 and RAB-27 are both enriched in the nerve ring of *C. elegans* (Mahoney et al. 2006), suggesting synaptic localization, and both Rabs colocalize at synapses in mammalian neurons (Pavlos et al. 2010). Consistent with these studies, we found that tagged *rab-3* and *rab-27* colocalize at synapses in *C. elegans* GABA neurons (Fig. 2A). *rab-3* regulates synaptic vesicle tethering and synaptic transmission (Mahoney et al. 2006), and *rab-27* is thought to play an auxiliary role in this process (Mahoney et al. 2006, Pavlos et al. 2010). Further, both *rab-27* and *rab-3* are regulated by a common GEF MADD/*aex-3*, and *aex-3* is required for normal synaptic transmission (Mahoney et al. 2006). However, despite these similarities, other data suggest that *rab-27* and *rab-3* also have different functions. In *C. elegans*, the Rab effector protein Rabhilin/*rbf-1* genetically interacts with *rab-27* but not *rab-3* (Mahoney et al. 2006, Mesa et al. 2011, Barclay et al. 2012). Further, *rab-27* and *rbf-1*, but not *rab-3*, are required for tethering and secretion of dense core vesicles in neurons (Ch’ng et al. 2008, Feng et al. 2012, Laurent et al. 2018). Finally, *rab-27*, unlike *rab-3* or Rabphilin/*rbf-1*, is expressed in both neurons and intestine (Mesa et al. 2011, Cao et al. 2017). Consistent with this, *rab-27* mutants but not *rab-3* or Rabphilin/*rbf-1* mutants have a constipated phenotype due to a defect in dense core vesicle release from the intestine and resulting disruption of the defecation motor program (DMP) (Riddle et al. 1997, Mahoney et al. 2008). These data raise the question of what the relationship is between *rab-27* and *rab-3* in axon regeneration.

**Figure 2.**
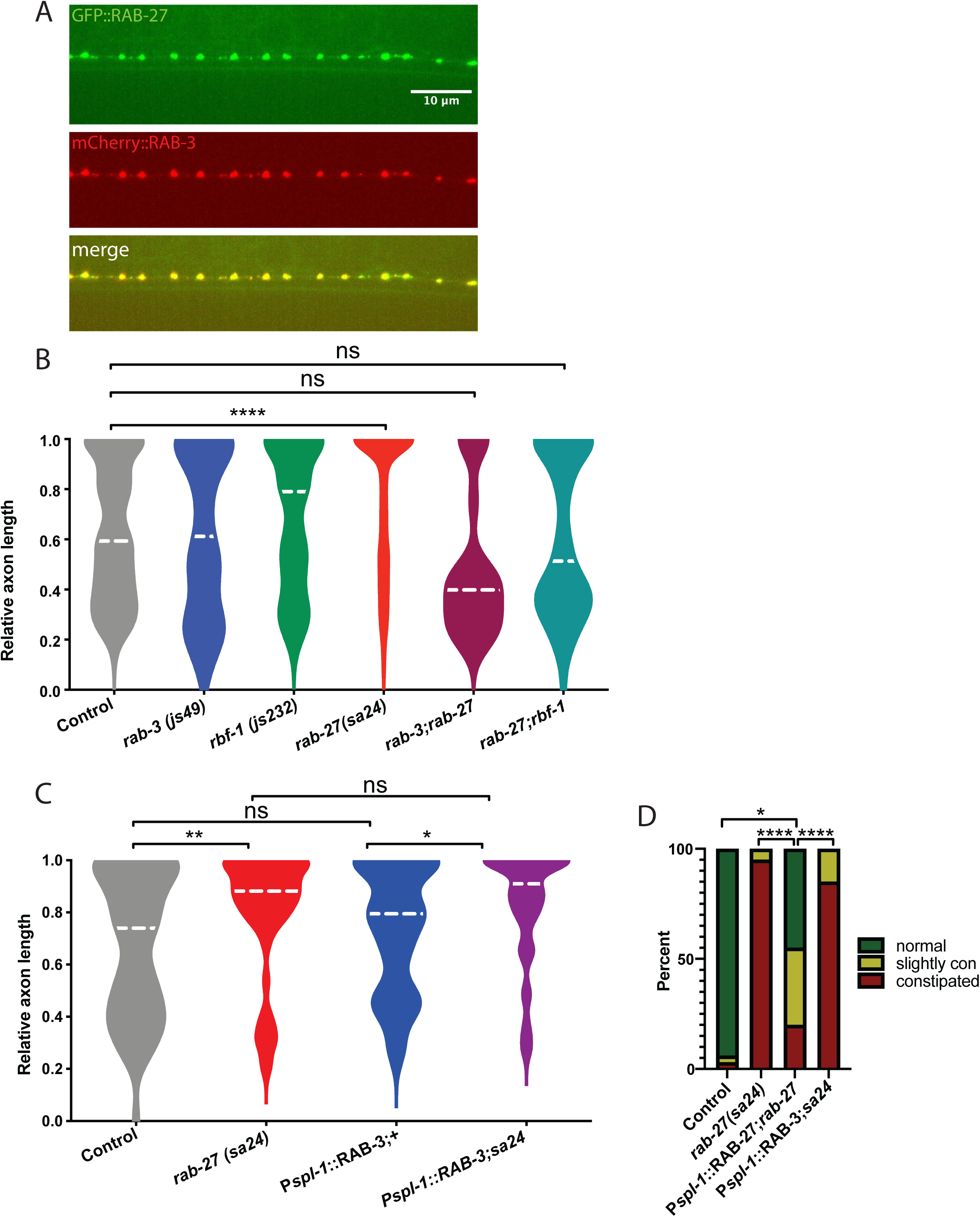
RAB-27’s synaptic vesicle tethering cofactors do not inhibit regeneration. (A) Colocalization of transgenic GFP::RAB-27 and mCherry::RAB-3 at synapses of DD/VD neurons. GFP::RAB-27 and mCherry::RAB-3 were expressed as multicopy arrays at an injection concentration of 7.5ng/μL. GFP::RAB-27 was expressed as multicopy array with a soluble mCherry transcriptional reporter at an injection concentration of 7.5ng/μL. (B) Relative axon length in control (*oxIs12*) animals, *rab-3(js49), rbr-1(js232), rab-27(sa24), rab-3(js49);rab-27(sa24)* mutants. Axons cut per genotype, L to R: 183, 37, 55, 196, 21, 69. Kolmogorov-Smirnov test was used. ns, not significant, * p < 0.05, *** p < 0.0005. (C) Relative axon length in control animals, *rab-27(sa24)* mutants, and animals expressing RAB-3 cDNA under an intestine-specific promoter, in control and *rab-27* mutant backgrounds. Number of axons cut per genotype, L to R: 61, 55, 53, 50. Kolmogorov-Smirnov test was used. ns, not significant, * p < 0.05, ** p < 0.005. (D) Percent stacked bar graph for visual scoring of Aex phenotype rescue. Animals were randomized on plates and scored by phenotype, then genotyped. Animals were scored as normal (no gut distention, strong pBoc contraction with accompanying expulsion), constipated (severe posterior gut distention, weak pBoc with no expulsion), or slightly con (some possible gut distention, normal pBoc, weak expulsion). Fisher’s Exact test was used. * p < 0.05, **** p < 0.0001. (E). Visualization of Aex phenotype and rescue in control and transgenic animals. Distention of the intestinal lumen, caused by failure to expel waste is characteristic of *rab-27* mutant animals, and was partially rescued by intestinal expression of RAB-27 cDNA, but not by RAB-3.

We used genetic analysis to determine the relationship between *rab-27, rab-3* and the effector Rabphilin/*rbf-1* in axon regeneration. Loss of *rab-3* did not affect axon regeneration (Fig. 2B). Thus, unlike for synaptic vesicle release, where *rab-3* predominates (Mahoney et al. 2006), *rab-27* rather than *rab-3* is the major factor in axon regeneration. Loss of Rabphilin/*rbf-1* also did not affect regeneration. However, double mutants for either *rab-27;rab-3* or *rab-27;rbf-1* suppressed the high regeneration phenotype of *rab-27* single mutants (Fig. 2B). We conclude that a neuronal function mediated by *rab-3* and Rabphilin/*rbf-1* is required for enhanced regeneration in *rab-27* mutants, though this neuronal function is dispensable for normal regeneration.

A major site of *rab-27* function in axon regeneration is the intestine (Fig. 1G), where *rab-3* is not expressed (Nonet et al. 1997). Given the close evolutionary and functional relationship between *rab-27* and *rab-3*, it is possible that *rab-3* could function in the intestine to inhibit axon regeneration, but is simply not expressed there. To test this idea, we ectopically expressed RAB-3 in the intestine of *rab-27* mutants to see whether RAB-3 could compensate for loss of *rab-27*. Intestinal expression of RAB-3 in *rab-27* mutants was not sufficient to rescue high regeneration (Fig. 2C). Intestinal RAB-3 also failed to rescue DMP defects in *rab-27* mutants. Thus, for the two distinct phenotypes of axon regeneration and DMP, *rab-27* mutants expressing intestinal RAB-3 were indistinguishable from non-transgenic *rab-27* mutants. By contrast, *rab-27* mutants expressing intestinal RAB-27 significantly rescued the DMP (Fig. 2D, Fig. S2), as well as restoring normal levels of axon regeneration (Fig. 2C). Together, these results indicate that despite their similarity and shared function in synaptic vesicle tethering, RAB-27 and RAB-3 are functionally distinct, and raise the question of what mechanisms act with RAB-27 to mediate its intestinal function in axon regeneration.

### Intestinal components of a secretory vesicle signaling pathway inhibit regeneration

In the intestine, *rab-27* acts to facilitate the tethering and fusion of dense core vesicles during the defecation motor program (DMP) (Mesa et al. 2011). At the expulsion (‘Exp’) step of the DMP, a neuropeptide ligand packaged into DCVs is secreted from the intestine. This peptide signal is sensed by receptors on the GABAergic neurons AVL and DVB, which in drive contractions of the enteric muscles and eventually waste expulsion (Riddle et al. 1997, Mahoney et al. 2008, Wang et al. 2013). Packaging and fusion of these intestinal DCVs involves *rab-27*, together with the pro-protein convertase KPC3/*aex-5*, the t-SNARE protein SNAP25/*aex-4*, the Munc13-like SNARE regulator *aex-1*, the Rab GEF recruitment factor NPDC1/*cab-1*, and the Rab GEF MADD/*aex-3*. The neuronal receptor that responds to neuropeptide release from the intestine is the GPCR *aex-2*. Loss of function in any of these genes disrupts the DMP and results in a constipation phenotype (Riddle et al. 1997, Mahoney et al. 2008, Wang et al. 2013).

We hypothesized that this same DCV secretion mechanism may account for *rab-27*’s function in axon regeneration. Consistent with this hypothesis, we found that KPC3/*aex-5*, SNAP25/*aex-4*, and NPDC1/*cab-1* all inhibit axon regeneration similar to *rab-27* mutants (Fig. 3B, Fig. 4A). However, loss of the Rab GEF MADD/*aex-3*, Munc13-b/*aex-1*, or the GPCR *aex-2* did not affect regeneration (Fig. 3B). Altogether, these results indicate that neuropeptide secretion from the intestine regulates axon regeneration, and that RAB-27 is an essential part of the secretion mechanism. However, this secretory pathway is genetically separable from the defecation motor program, suggesting that regulation of axon regeneration involves a distinct, specialized pool of DCVs.

**Figure 3.**
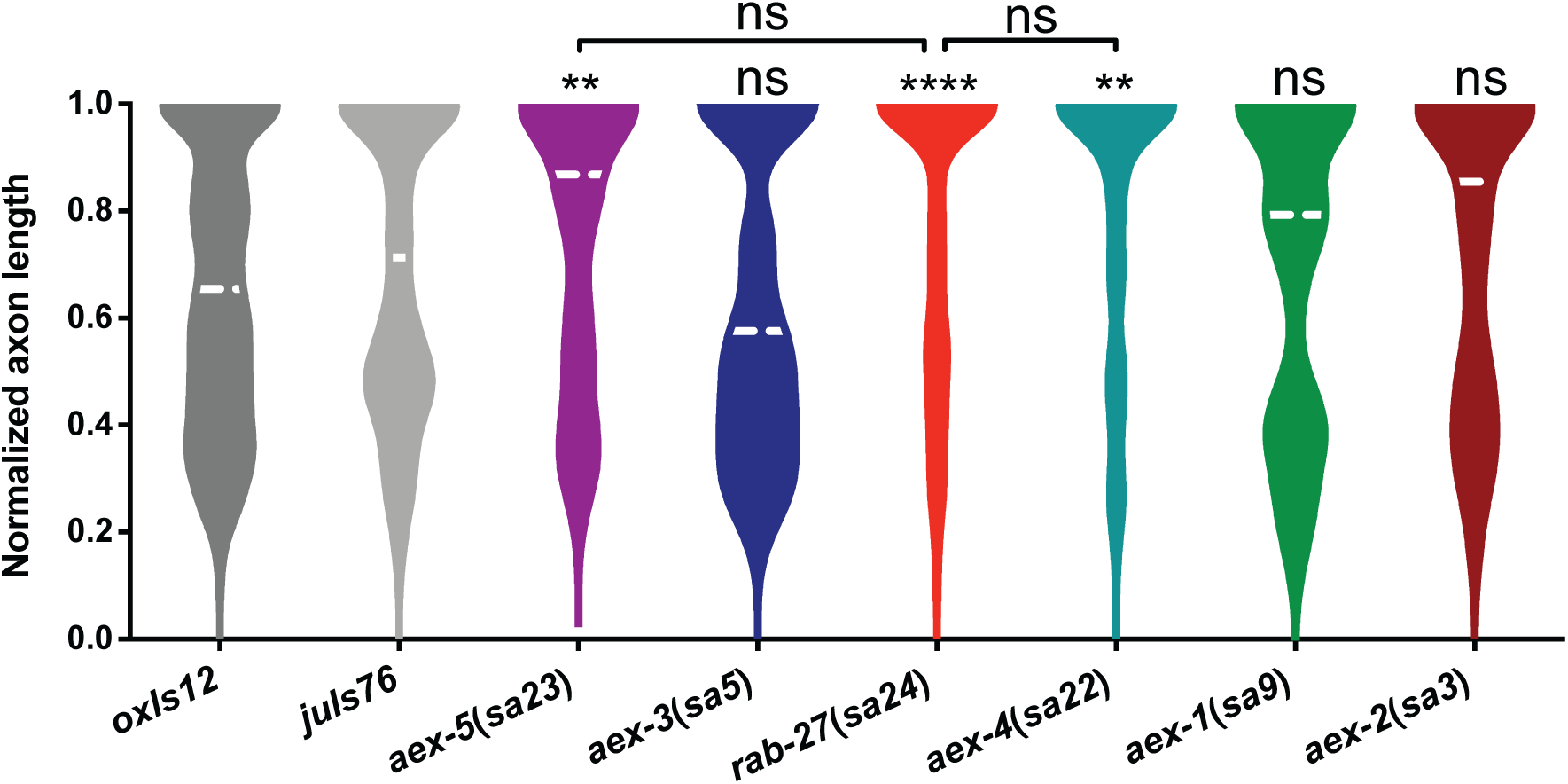
AEX-4 and AEX-5 inhibit axon regeneration. (A) Relative axon length in control animals expressing GABAergic neuron-specific GFP (*oxIs12* & *juIs76*), and *aex-1(sa9), aex-2(sa3), aex-3(sa5), aex-4(sa22), aex-5(s23)* and *rab-27(sa24)* mutants. *aex-1, aex-5*, and *rab-27* are compared against *oxIs12*, while *aex-2, aex-3*, while *aex-4* are compared against *juIs76*. Axons cut per genotype, L to R: 238, 199, 37, 83, 148, 69, 50, 66. Kolmogorov-Smirnov test was used. ns, not significant, * p < 0.05, ** p < 0.005 **** p < 0.0001.

**Figure 4.**
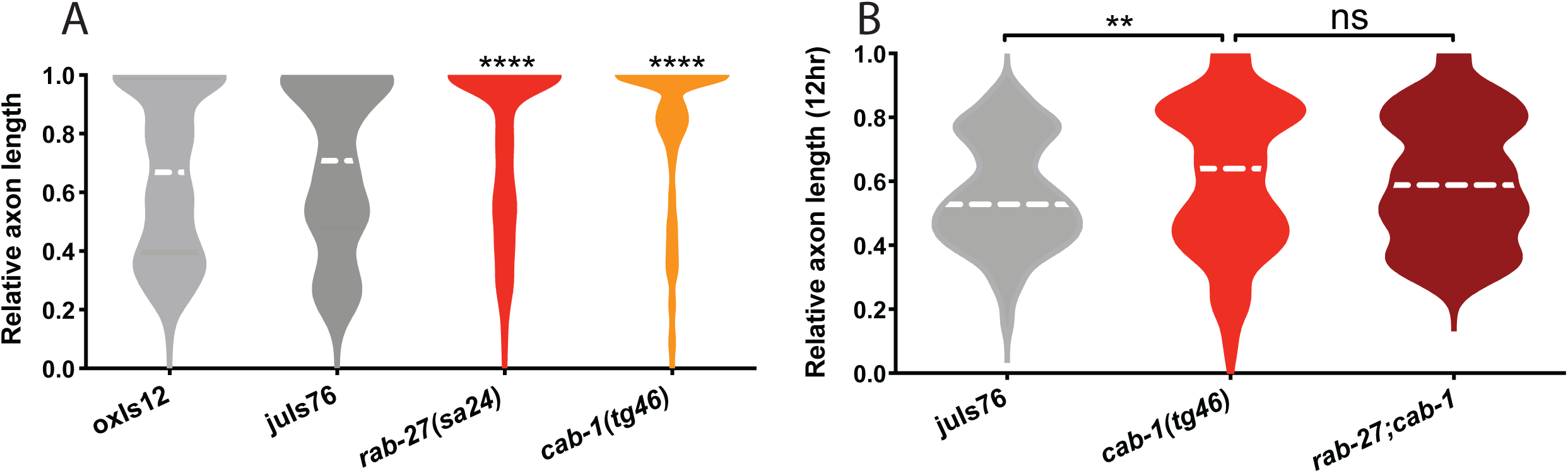
CAB-1 inhibits axon regeneration. (A) Relative axon length in control animals expressing GABAergic neuron-specific GFP (*oxIs12* & *juIs76*), and *rab-27(sa24)* and *cab-1(tg46)* mutants. *rab-27* is compared against *oxIs12*, while *cab-1* is compared against *juIs76*. Axons cut per genotype, L to R: 200, 81, 164, 91. Kolmogorov-Smirnov test was used. ns, not significant, **** p < 0.0001. (B) Relative axon length in control animals expressing GABAergic neuron-specific GFP (*juIs76*), *rab-27(sa24)* mutants and *rab-27(sa24);cab-1(tg46)* double mutants. L to R: 78, 64, 90. Kolmogorov-Smirnov test was used. ns, not significant, ** p < 0.005. Regeneration was scored after 12 hours of recovery to more easily visualize enhanced regeneration in the *rab-27* and *rab-27;cab-1* double mutants, which show nearly full regeneration after the usual 24 hour recovery window.

The identity of the secreted signal, and the receptors that transduce it, are presently unknown. Over 250 distinct neuropeptides have been identified in *C. elegans* (Li & Kim 2008), of which approximately fifty are believed to be expressed in the intestine (Nathoo et al. 2001, Pierce et al. 2001, Li et al. 2003, Cao et al. 2017). A small candidate screen of intestinally-expressed neuropeptide-like proteins (NLPs) that are expressed in the intestine and are processing targets of KPC3/AEX-5 (Husson et al. 2006) did not identify any inhibitors of regeneration (Fig. S5). Similarly, the *C. elegans* has between 125 to 150 G-protein coupled neuropeptide receptor homologs (Frooninckx et al. 2012, Koelle 2018), of which approximately 20 are expressed in the DD/VD GABAergic motor neurons (Taylor et al. 2019). Of these, we find that the GPCR AEX-2 does not inhibit regeneration, although it does respond to peptide signals from the intestine in the context of the DMP (Wang et al. 2013). The identity of the peptide signal or signals, and the potential receptor remain unknown. Further work is required to identify these components of the intestine-neuron signaling axis that inhibits axon regeneration.

### Multiple Rab GTPases affect axon regeneration

*rab-27* was initially identified as a candidate regeneration inhibitor in a functional genome-wide screen for regeneration inhibitors done in mammalian cortical neurons *in vitro* that identified 19 Rab GTPases as potential regeneration inhibitors (Sekine et al. 2018). *C. elegans* has a drastically reduced cohort of functional Rabs compared to mammals (Gallegos et al. 2012), attributable in large part to decreases in redundancy. Compared to the results seen in mammalian cell culture, a few Rabs in *C. elegans* affect regeneration (Fig. 5A). In addition to *rab-27* and the previously identified *rab-6*.*2* (Zeng et al. 2018), loss of *rab-18* significantly decreases regeneration success, while loss of *glo-1* leads to a modest increase in regeneration. Unlike other high-regenerating Rab mutants, *glo-1* mutants specifically show an increase in full regeneration after 24 hours of recovery, though not an increase in the likelihood of regeneration initiation during that period (Fig. 5B,C). GLO-1 is expressed specifically in the intestine, where it localizes to and is required for the biogenesis of the lysosome-like gut granules (Hermann et al. 2005). Along with *rab-27*, the effect of *glo-1* on regeneration suggests that the intestine may play a previously unknown but important role in regulation of axon regeneration.

**Figure 5.**
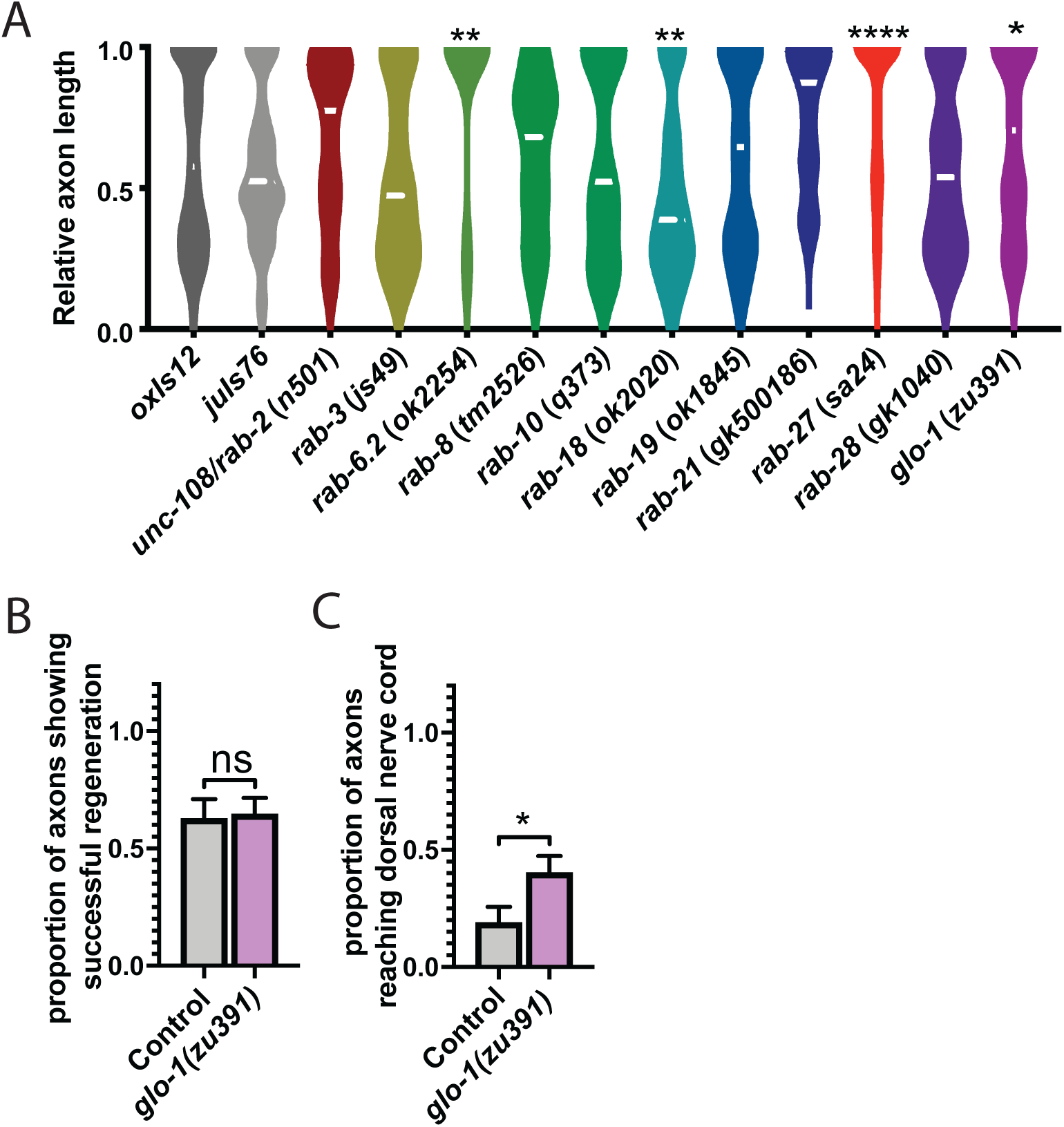
Multiple Rab GTPases affect axon regeneration. (A) (B) Relative axon length in control animals expressing GABAergic neuron-specific GFP (*oxIs12* & *juIs76*), and *unc-108/rab-2(n501), rab-3(js49), rab-6*.*2(ok2254), rab-8(tm2526), rab-10(q373), rab-18(ok2020), rab-19(ok1845), rab-21(gk500186), rab-27(sa24), rab-28(gk1040), and glo-1(zu391). unc-108/rab-2, rab-3, rab-8, rab-10, rab-18, rab-19, rab-21, rab-27* and *rab-28* are compared against *oxIs12*, while *rab-6*.*2* and *glo-1* are compared against *juIs76*. Axons cut per genotype, L to R: 396, 46, 39, 72, 13, 25, 41, 69, 43, 38, 123, 21, 45, 64. Kolmogorov-Smirnov test was used. ns, not significant, * p < 0.05, ** p < 0.005 **** p < 0.0001. (B) Proportion of cut axons showing signs of regeneration in control (*juIs76*) and *glo-1(zu391)* mutant animals. Axons cut per genotype, L to R: 32, 45. Unpaired t-test was used. ns, not significant. Error bars represent SEM. (C) Proportion of cut axons showing full regeneration back to the dorsal nerve cord in control (*juIs76*) and *glo-1(zu391)* mutant animals. Axons cut per genotype, L to R: 32, 45. Unpaired t-test was used. ns, not significant. Error bars represent SEM.

## DISCUSSION

Axon regeneration is tightly regulated by pathways from within the injured neuron as well as by interactions with the local environment, but the existence of long-range regulatory signals has remained unclear. Here we show that in *C. elegans*, RAB-27 acts in the intestine to inhibit regeneration of severed axons of the DD/VD GABAergic motor neurons. This inhibition occurs independently of *rab-27*’s known role in neurons, where it regulates synaptic vesicle fusion and also functions in axon regeneration (Mahoney et al. 2006, Sekine et al. 2018).

We find that multiple factors involved in dense core vesicle (DCV) packaging and secretion from the intestine inhibit regeneration along with *rab-27*. Specifically, CAB-1 and SNAP25/AEX-4, which function in DCV trafficking and fusion (E. Jorgensen, pers. comm., Mahoney et al. 2008, Xia et al. 2014,), and KPC3/AEX-5 which functions in neuropeptide processing (Husson et al. 2006), inhibit regeneration. These data suggest a model in which axon regeneration is regulated by a neuropeptide signal, processed by KPC3/AEX-5, that is packaged into dense core vesicles, tether to the basal membrane of intestinal cells via RAB-27-dependent interactions, and secreted via SNAP25/AEX-4-dependent SNARE activity. An attractive hypothesis is that a neuronal neuropeptide receptor responds to this signal to limit regeneration.

Surprisingly we find no role for Munc-13b/*aex-1* in regeneration. Munc13 proteins are involved in SNARE-mediated vesicle docking and fusion (Hammarlund et al. 2007, Lai et al. 2017), and Munc13-b/*aex-1* is required for DCV fusion in the intestine during the DMP (Yamashita et al. 2009). These data suggest that the intestinal DCV population that mediates regeneration is distinct from DCVs that mediate the DMP. Presumably the “regeneration DCVs” rely on a different factor than the “DMP DCVs” to mediate SNARE-directed fusion. However, we did not detect a role in regeneration for CAPS/*unc-31*(Fig. S3), another factor that mediates SNARE-directed membrane fusion (Hammarlund et al. 2008). One possibility is that Munc-13b/AEX-1 may function redundantly with other vesicle docking regulators to mediate DCV fusion for axon regeneration.

In the nervous system, RAB-27 regulates synaptic vesicle tethering in coordination with the closely related RAB-3, upstream of the effector Rabphilin/RBF-1 (Mahoney et al. 2006, Mesa et al. 2011). While neuronal RAB-27 inhibits regeneration (Fig. 1H), loss of *rab-3* or Rabphilin/*rbf-1* does not affect regeneration (Fig. 2B). These data suggest that neuronal RAB-27 inhibits axon regeneration independent of its role in synaptic vesicle tethering. As it does in diverse tissues across species, RAB-27 also regulates the tethering and fusion of non-synaptic vesicles in *C. elegans* neurons (Feng et al. 2012), and similar to the intestine, neuronal RAB-27 may regulate the secretion of an unknown ligand or ligands through dense core vesicles to inhibit regeneration. Several possibilities could explain neuronal RAB-27’s incomplete rescue of high regeneration compared to intestinal RAB-27: the two tissue-specific RAB-27-dependent pathways may be regulating the release of different inhibitory ligands, with the intestine secreting a more potent inhibitor. Alternatively, intestinal and neuronal RAB-27 could be promoting release of the same inhibitory ligand or ligands, with these ligands highly secreted from the intestine but only marginally expressed in neurons.

While loss of *rab-3* or Rabphilin/*rbf-1* alone does not affect regeneration, loss of either in a *rab-27* mutant background completely suppresses the *rab-27* mutant high regeneration phenotype (Fig. 2B). However, these double mutants, which show severe defects in synaptic transmission (Mahoney et al. 2006), do not show any defects in regeneration beyond the suppression of the *rab-27* mutant phenotype (Fig 2B). These data suggest that robust synaptic vesicle fusion is required only for enhanced regeneration. Significant loss of vesicle fusion below a certain threshold may restrict high regeneration by restricting the available pool of membrane required for enhanced outgrowth (Futerman & Banker 1996). Alternatively, loss of synaptic vesicle tethering and fusion could disrupt specific pro-regeneration pathways that are normally inhibited during regeneration, but that are released following loss of inhibitory upstream regulatory signals such as RAB-27. Thus, neuronal RAB-27 appears to have dual roles in the regulation of axon regeneration: a pro-high regenerative role mediated through synaptic vesicle fusion and co-regulated by RAB-3 and Rabphilin/RBF-1, and an inhibitory role mediated by the secretion of an anti-regeneration signal from DCV fusion.

Rab GTPases are emerging as key regulators of axon regeneration *in vitro* and *in vivo. C. elegans* provides an excellent system to probe the “rabome” for novel pathways affecting axon regeneration. In *C. elegans, rab-6*.*2* was previously shown to affect regeneration (Zeng et al. 2018), as was *rab-27* function in neurons (Sekine et al. 2018). This work probed the function of RAB-27 outside the nervous system, revealing an unexpected role for DCV fusion in the intestine in regulation of axon regeneration. Rabs mediate many complex biological processes, such as Parkinson’s disease pathogenesis (Gao et al. 2018) and cancer metastasis through regulation of exosome secretion (Li et al. 2018). This study adds to our understanding of Rab function by identifying a novel role for RAB-27 in mediating a long-range signal that inhibits the ability of neurons to regenerate after injury.

## Materials and Methods

### *C. elegans* strains

Strains were maintained at 20C, as described in Brenner (Brenner, 1974), on NGM plates seeded with OP50. Some strains were provided by the CGC, which is funded by the NIH Office of Research Infrastructure Programs (P40 OD010440). The following strains were purchased from the CGC:

NM791[*rab-3(js49)*], RT2[*rab-10(e1747)*], RB1638[*rab-18(ok2020*], RB1537[*rab-19(ok1845*], JT24[*rab-27(sa24)*], JT699[*rab-27(sa699)*], JJ1271[*glo-1(zu391)*], VC2505[*rab-28(gk1040)*], MT1093[*unc-108(n501)*], JT23[*aex-5(sa23)*], JT3[*aex-2(sa3)*], JT5[*aex-3(sa5)*], JT9[*aex-1(sa9)*], KY46[*cab-1(tg46)*], NM1278[*rbf-1(js232)*], NM2777 [*aex-6(sa24);rab-3(js49)*]. The following strains were purchased from the NBRP: *rab-8(tm2526)*.

List of generated strains:

|  |  |
| --- | --- |
| *rab-27(sa24)* I; *oxIs12[Punc-47::GFP;lin-15+]* X | XE1873 |
| wpEx434[*Pspl-1::RAB-27::SL2::mCherry::RAB-3 3' UTR*]; *oxIs12[Punc-47::GFP;lin-15+]* X | XE2524 |
| wpEx417[*Pspl-1::RAB-27::SL2::mCherry::RAB-3 UTR*];*rab-27(sa24)* I; *oxIs12[Punc-47::GFP;lin-15+]* X | XE2452 |
| wpEx418[*Punc-47::RAB-27::SL2::mCherry::RAB-3 3' UTR*]; *oxIs12[Punc-47::GFP;lin-15+]* X | XE2451 |
| wpEx436[*Punc-47::RAB-27::SL2::mCherry::RAB-3 3' UTR*];*rab-27(sa24)* I; *oxIs12[Punc-47::GFP;lin-15+]* X | XE2525 |
| wpEx287[*Punc-47::RAB-27::SL2::mCherry::UNC-54 3' UTR*]; *oxIs12[Punc-47::GFP;lin-15+]* X | XE1874 |
| wpEx287[*Punc-47::RAB-27::SL2::mCherry::UNC-54 3' UTR*];*rab-27(sa24)* I; *oxIs12[Punc-47::GFP;lin-15+]* X | XE1890 |
| wpEx405[*Pspl-1::RAB-27::SL2::mCherry::UNC-54 3' UTR*]; *oxIs12[Punc-47::GFP;lin-15+]* X | XE2353 |
| wpEx405[*Pspl-1::RAB-27::SL2::mCherry::UNC-54 3' UTR*];*rab-27(sa24)* I; *oxIs12[Punc-47::GFP;lin-15+]* X | XE2354 |
| wpEx288[*Punc-47::EGFP::RAB-27::UNC-54 3' UTR*]; *wpls40[Punc-47::mCherry]* V | XE1904 |
| wpEx435[*Punc-47::EGFP::RAB-27::UNC-54 3' UTR*; *Punc-47::mCherry::RAB-3::UNC-54 3' UTR*] | XE2523 |
| *rab-3(js49)* II; *oxIs12[Punc-47::GFP;lin-15+]* X | XE1871 |
| *rbf-1(js232)* III; *oxIs12[Punc-47::GFP;lin-15+]* X | XE1901 |
| *rab-27(sa24)* I; *rab-3(js49)* II; *oxIs12[Punc-47::GFP;lin-15+]* X | XE2514 |
| *rab-27(sa24)* I; *rbf-1(js232)* III; *oxIs12[Punc-47::GFP;lin-15+]* X | XE2515 |
| wpEx406[*Pspl-1::RAB-3::SL2::mCherry::UNC-54 3' UTR*];*rab-27(sa24)* I; *oxIs12[Punc-47::GFP;lin-15+]* X | XE2351 |
| wpEx406[*Pspl-1::RAB-3::SL2::mCherry::UNC-54 3' UTR*]; *oxIs12[Punc-47::GFP;lin-15+]* X | XE2352 |
| *aex-1(sa9)* I; *oxls12[Punc-47::GFP;lin-15+]* X | XE2511 |
| *aex-2(sa3)* X; *juls76[Punc-25::GFP;lin-15+]* II | XE2517 |
| *aex-3(sa5)* X; *juls76[Punc-25::GFP;lin-15+]* II | XE2510 |
| *aex-4(sa22)* X; *juls76[Punc-25::GFP;lin-15+]* II | XE2516 |
| *aex-5(sa23)*; *oxls12[Punc-47::GFP;lin-15+]* X | XE2509 |
| *unc-31(e928)* IV; *oxls12[Punc-47::GFP;lin-15+]* X | XE1905 |
| *cab-1(tg46)* X; *juls76[Punc-25::GFP;lin-15+]* II | XE2512 |
| *cab-1(tg46)* X; *rab-27(sa24)* I; *juls76[Punc-25::GFP;lin-15+]* II | XE2513 |
| *unc-108/rab-2(n501)* I; *oxls12[Punc-47::GFP;lin-15+]* X | XE2518 |
| *rab-6.2(ok2254)* X; *juls76[Punc-25::GFP;lin-15+]* II | XE1560 |
| *rab-8(tm2526)* I; *oxls12[Punc-47::GFP;lin-15+]* X | XE2519 |
| *rab-10(q373)* I; *oxls12[Punc-47::GFP;lin-15+]* X | XE1804 |
| *rab-18(ok2020)* III; *oxls12[Punc-47::GFP;lin-15+]* X | XE1872 |
| *rab-19(ok1845)* IV; *oxls12[Punc-47::GFP;lin-15+]* X | XE2522 |
| *rab-21(gk500186)* II; *oxls12[Punc-47::GFP;lin-15+]* X | XE2521 |
| *rab-28(gk1040)* IV; *oxls12[Punc-47::GFP;lin-15+]* X | XE1806 |
| *glo-1(zu391)* X; *juls76[Punc-25::GFP;lin-15+]* II | XE2520 |
| *nlp-1(ok1469)* X; *juls76[Punc-25::GFP;lin-15+]* II | XE2409 |
| *nlp-8(ok1799)* I; *juls76[Punc-25::GFP;lin-15+]* II | XE2407 |
| *nlp-20(ok1591)* IV; *juls76[Punc-25::GFP;lin-15+]* II | XE2408 |

### Constructs and cloning

Transgenic constructs were generated with Gateway recombination (Invitrogen). Fluorescent-tagged RAB-27 was generated through fusion PCR (Hobert 2002)

### Laser axotomy

Laser axotomy was performed as previously described in Byrne et al. 2011. L4 animals were immobilized using 0.05 µm polystyrene beads (Polybead Microspheres, Polysciences Cat #08691-10) or in 0.2mM Levamisole (Sigma) on a pad of 3% agarose dissolved in M9 buffer on a glass slide. Worms were visualized using a Nikon Eclipse 80i microscope with a 100x Plan Apo VC lens (1.4 NA). Fluorescently-labeled D-type motor neuron commissures were targeted at the dorsoventral midline using a 435 nm Micropoint laser with 10 pulses at 20 Hz. In all cases no more than four of the seven posterior commisures were cut per animal to minimize possible adverse locomotion or behavioral effects. Animals were recovered to NGM plates seeded with OP50 and allowed to recover.

### Fluorescence microscopy and regeneration scoring

Animals with cut axons were immobilized using 0.25–2.5 mM levamisole (Santa Cruz, sc-205730) and mounted on a pad of 3% agarose in M9 on glass slides. All animals were imaged to visualize regeneration using an Olympus DSU mounted on an Olympus BX61 microscope, with a Hamamatsu ORCA-Flash4.0 LT camera, and Xcite XLED1 light source with BDX, GYX and RLX LED modules. Images were acquired as 0.6 um z-stacks using consistent exposure time, camera sensitivity and light intensity. Images were exported as tiff files and analyzed in ImageJ. Cut axons were scored based on regeneration status and length, and each individual axon was given a designation showing presence of a growth cone indicative of regeneration initiation (Y,N), its general elongation status (no regeneration, GC below midline, GC at midline, GC above midline, full regeneration to DNC), and the measured axon length (absolute axon growth relative to the distance between dorsal and ventral nerve cords). Significance is indicated by and asterisk (*p<0.05, Kolmogorov-Smirnov test).

For imaging of GFP::RAB-27 in cut axons (Fig. S1C-E) and GFP::RAB-27; mCherry::RAB-3 in intact axons (Fig. 2A), worms were immobilized as described above, and imaged using the vt-iSIM system mounted on a Leica DMi8 inverted platform, with a Hammamatsu ORCA-Flash 4.0 camera. Images were acquired as 0.6 um z-stacks using consistent exposure time, camera sensitivity and light intensity.

### Fecundity

L4 worms of each genotype were singled onto NGM plates seeded with 100µL OP50 for 48 hours. Adult worms were removed, and surviving progeny (L1 or older animals) were counted after an additional 24 hours. Unhatched eggs were not counted.

## Supporting information

Figure S2. Rescue of the defecation motor program by intestinal rab-27 expression

Figure S3. Two dense core vesicle tethering regulators do not affect axon regeneration

Figure S4. cab-1 and rab-27 show reduced fecundity

Figure S5. Loss of nlp-1, nlp-8 or nlp-20 does not affect axon regeneration

## Acknowledgements

We thank WormBase and the Caenorhabditis Genetics Center (CGC), which is funded by the National Institutes of Health (NIH) Office of Research Infrastructure Programs (P40 OD010440). We also thank Tyler Page and Erik Jorgensen for suggestions and feedback regarding *cab-1*. This research was supported by NIH grants (R01 NS098817 and R01 NS094219) to M.H.

## Author contributions

A.L.M. and M.H. designed experiments. A.L.M. and R.J.O. performed experiments and data analysis.

