## Supplementary figures and images for "*rab-27* acts in an intestinal secretory pathway to inhibit axon regeneration in *C. elegans*"

### Figure S2. Rescue of the defecation motor program by intestinal rab-27 expression

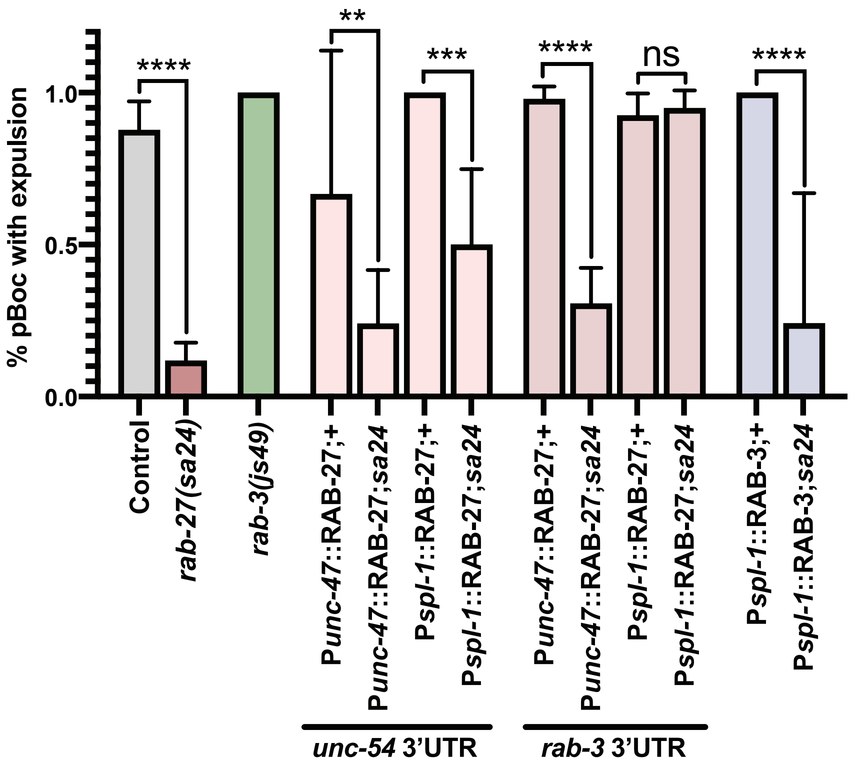

### Figure S3. Two dense core vesicle tethering regulators do not affect axon regeneration

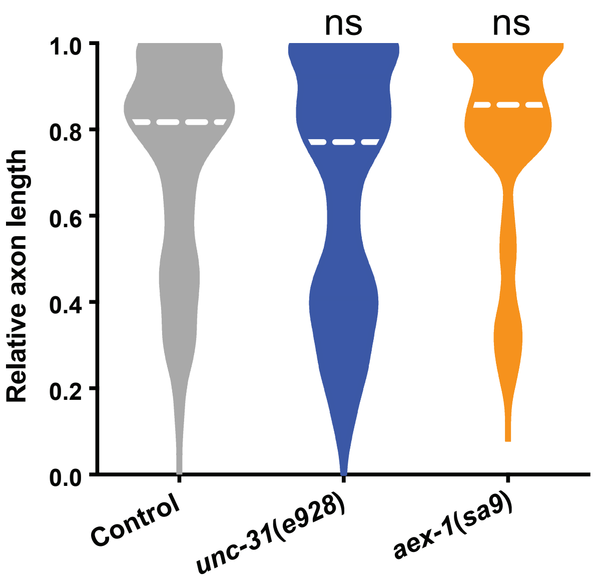

### Figure S4. cab-1 and rab-27 show reduced fecundity

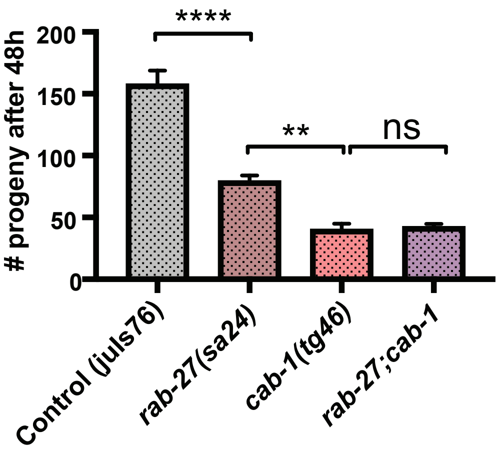

### Figure S5. Loss of nlp-1, nlp-8 or nlp-20 does not affect axon regeneration

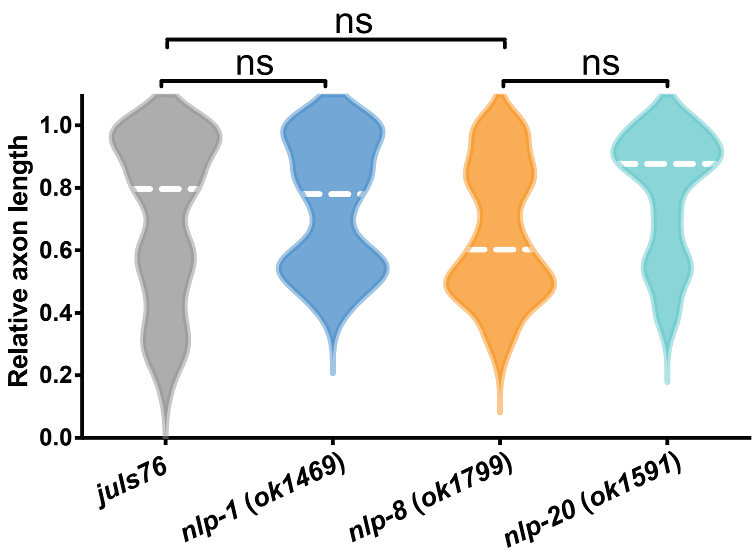

### Supplemental Data 1

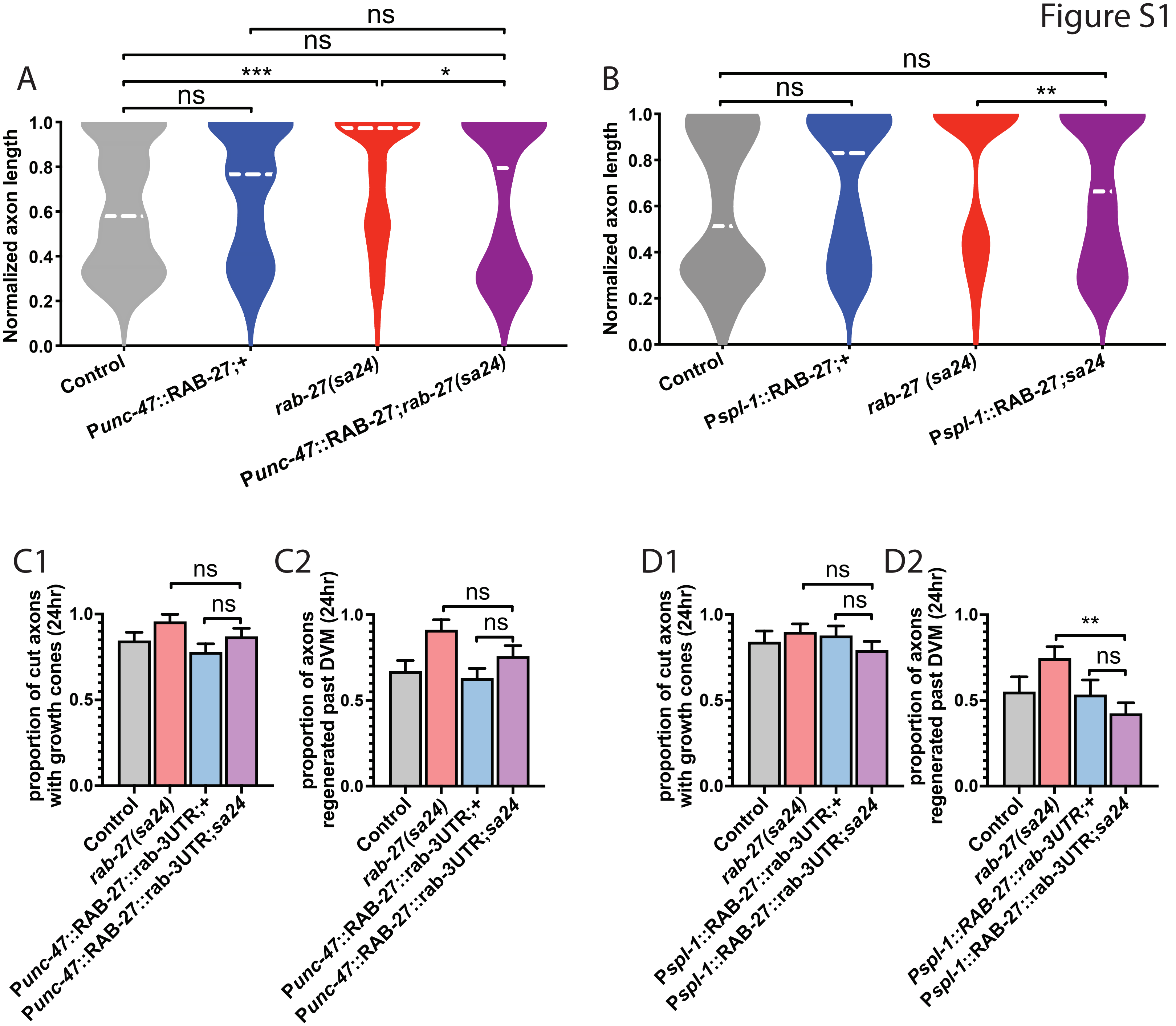
