## Supplemental figure legends for "*rab-27* acts in an intestinal secretory pathway to inhibit axon regeneration in *C. elegans*"

**Figure S1**. Use of *unc-54* 3’ UTR sequence in constructs containing RAB-27 cDNA inhibits regeneration. (A-B) Relative axon length in animals expressing RAB-27 cDNA under a GABA neuron-specific (A) or intestine-specific (B) promoter and with *unc-54* 3’ UTR sequence, in both control (*oxIs12*) and *rab-27* mutant backgrounds. Number of axons cut per genotype, L to R: 51, 67, 22, 45. Kolmogorov-Smirnov test was used. ns, not significant, * p < 0.05, ** p < 0.005, *** p < 0.0005. (C) Proportion of cut axons showing signs of successful regeneration initiation (C1) or regeneration past the dorsoventral midline (C2) in control (*oxIs12*) and *rab-27(sa24)* mutant animals, and animals expressing *rab-27* cDNA under a GABA neuron-specific promoter (Punc-47) and the *rab-3* 3’ UTR sequence, in both control and *rab-27* mutant backgrounds. Axons were scored after 24 hours of recovery post-axotomy. Axons cut per genotype, L to R: 51, 22, 67, 45. Unpaired t-test was used. ns, not significant. Error bars represent SEM. (D) Proportion of cut axons showing signs of successful regeneration initiation (D1) or regeneration past the dorsoventral midline (D2) in control (*oxIs12*) and *rab-27(sa24)* mutant animals, and animals expressing *rab-27* cDNA under an intestine-specific promoter (Pspl-1) and the *rab-3* 3’ UTR sequence, in both control and *rab-27* mutant backgrounds. Axons were scored after 24 hours of recovery post-axotomy. Axons cut per genotype, L to R: 31, 39, 32, 57. Unpaired t-test was used. ns, not significant, ** p < 0.005. Error bars represent SEM.

**Figure S2**. Rescue of the defecation motor program by intestinal *rab-27* expression. Mutants in the *aex* pathway display a defect in the defecation motor program, visualized by a loss of waste expulsion (Exp) following posterior body contraction (pBoc). Animals were randomly selected and observed for 5 DMP cycles, and the ratio of Exp/pBoc was plotted. Intestinal (P*spl-1*) but not GABA neuron-specific (P*unc-47*) expression of *rab-27* cDNA was sufficient to rescue DMP in *rab-27* mutant worms. This rescue was enhanced in animals expressing constructs with a *rab-3* 3’ UTR compared to animals expressing constructs with a *unc-54* 3’ UTR. Expression of *rab-3* cDNA in the intestine of *rab-27* mutant animals did not rescue DMP defects. pBoc cycles observed, L to R: 49, 119, 30, 27, 25, 20, 18, 49, 62, 54, 56, 40, 58. Kolmogorov-Smirnov test was used. ns, not significant, * p < 0.05, ** p < 0.05, *** p < 0.0005, **** p < 0.0001, Fisher’s Exact Test. Error bars represent SEM.

**Figure S3**. Two dense core vesicle tethering regulators do not affect axon regeneration. (A) Relative axon length in control (*juIs76*) animals, *unc-31(e928)* and *aex-1(sa9)* mutants. Axons cut per genotype, L to R: 91, 59, 116. Kolmogorov-Smirnov test was used. ns, not significant.

**Figure S4**. *cab-1* and *rab-27* show reduced fecundity. One-day adult worms were placed onto empty NGM plates seeded with OP50 and left for 48 hours. Adults were removed and progeny counted. *rab-27* mutants show significantly decreased brood size compared to control animals, and *cab-1* mutants show more severe defects. The low brood size of *cab-1* mutants is not increased in *rab-27;cab-1* double mutants. Worms sampled, L to R: 9, 10, 7, 8. One-way ANOVA test was used. ns, not significant, ** p < 0.005, **** p < 0.0001. Error bars represent SEM.

**Figure S5**. Loss of *nlp-1, nlp-8* or *nlp-20* does not affect axon regeneration. Relative axon length in control (*juIs76*) animals, *nlp-1*(*ok1469*), *nlp-8*(*ok1799*), and *nlp-20*(*ok1591*) mutants. Axons cut per genotype, L to R: 58, 17, 47, 22. Kolmogorov-Smirnov test was used. ns, not significant.
